# Longitudinal high-resolution imaging through a flexible intravital imaging window

**DOI:** 10.1101/2021.02.24.432491

**Authors:** Guillaume Jacquemin, Maria Benavente-Diaz, Samir Djaber, Aurélien Bore, Virginie Dangles-Marie, Didier Surdez, Shahragim Tajbakhsh, Silvia Fre, Bethan Lloyd-Lewis

## Abstract

Intravital microscopy (IVM) is a powerful technique that enables imaging of internal tissues at (sub)cellular resolutions in living animals. Here, we present a silicone-based imaging window consisting of a fully flexible, suture-less design that is ideally suited for long-term, longitudinal IVM of growing tissues and tumors. Crucially, we show that this window, without any customization, is suitable for numerous anatomical locations in mice using a rapid and standardized implantation procedure. This low-cost device represents a substantial technological and performance advance that facilitates intravital imaging in diverse contexts in higher organisms, opening new avenues for *in vivo* imaging of soft and fragile tissues.

**One-sentence summary:** This study presents a versatile, fully flexible imaging window that acts as an implantable transparent ‘second skin’ for small laboratory animal *in vivo* imaging.

## Introduction

Bioimaging across multiple scales is a universal and mainstay tool in life science research. Technological advances in live and deep-tissue imaging methodologies, including intravital microscopy (IVM), now enable the real-time microscopic imaging of individual cells within intact tissues in near physiological conditions (*1*–*3*). This powerful approach is increasingly leveraged in experimental and pre-clinical studies to reveal novel insights into the dynamic cellular mechanisms underlying disease development and response to therapy (*4*, *5*). To facilitate repeated IVM over prolonged time periods in the same living animal, multiple imaging windows have been designed to provide optical access to internal tissues (*6*), including the brain (*7*), skin (*8*, *9*), lung (*10*), mammary gland (*6*, *11*, *12*), abdominal organs (*13*–*15*), femur (*16*) and embryos (*17*). Typically, these models consist of a glass coverslip inserted in a titanium (or more rarely plastic) frame and rely on sutures and/or glue to fix the window in place. Although powerful, this implantation method - combined with the material composition and rigidity of conventional windows - is poorly suited to dynamic or rapidly growing soft tissues or tumors, often requiring customization to meet tissue- or study-specific needs. The inherent rigidity of glass also limits its ability to adequately cover anatomical locations that exhibit curvature or joints. Collectively, these issues can cause animal distress, skin and tissue degradation, inflammation, fibrosis and window detachment, ultimately leading to experimental failures. Moreover, the reliance of all conventional windows on suturing (and occasionally glue) renders their implantation complicated and time-consuming. Finally, the absence of standardized and affordable imaging windows forces laboratories to make and reuse their own models, further precluding the universal and consistent application of IVM.

To address these limitations, here we developed a flexible and suture-less polydimethylsiloxane (PDMS)-based intravital imaging window dedicated to efficient, long-term maintenance at all body sites and improved animal welfare. While a few studies have reported the use of fixed-skull silicone membranes for brain IVM (*18*–*20*), the application of PDMS for soft tissue imaging remains unexplored, being limited to window designs that maintain the use of glass coverslips and sutures (*21*). As such, these models provide no benefit over conventional rigid windows. By contrast, our design encompasses a flexible, seamlessly-joined PDMS window that ensures a sealed barrier between the animal’s internal tissues and the external environment in a suture-free manner. By thorough characterization of its optical properties, we validate the utility of the PDMS membrane as an optical material for deep-tissue IVM. Alongside, we demonstrate the compatibility of the flexible window with intravital imaging at dynamic body locations and in conditions of extreme tissue growth, contexts that are less accessible to existing rigid windows. In turn, our engineered device promises to open new perspectives for *in vivo* imaging of soft and fragile tissues in higher vertebrates.

## Results

### PDMS-based intravital imaging window design

To address the restraints of current rigid models, we designed a flexible intravital imaging window made entirely of polydimethylsiloxane (PDMS) (Fig. 1A-B). PDMS is a light, biocompatible, chemically inert and optically clear material that is commonly used in medical or micro-fluidic devices. Importantly, it can be easily and reproducibly cast into any desired shape, size, thickness and rigidity by varying mold footprints, silicone composition and surface treatments. Collectively, these properties make PDMS an optimal material for devices demanding specifications for biocompatibility, versatility and optical performance. Our PDMS-based intravital imaging window comprises a supple and lightweight frame surrounding a thinner (145 μm ±10μm) area for imaging (Fig. 1B). A priority for the design of a universally adaptable window centered on a rapid and stereotyped suture-free implantation. To achieve this, we designed a groove around the window circumference that is precisely angled to secure the skin in place in the absence of sutures or glue (Fig.1B [1]). Passive sealing at the silicone-skin interface maintains the window post-implantation, while a series of holes around the frame promotes tissue self-healing for long term maintenance (Fig. 1B [2]). In addition to securing the skin inside the groove, this design maintains the window in a planar and near-seamless position in line with the animal’s body. Windows are therefore poorly accessible to the mouse and surrounding cage enrichment, reducing the risk of dislodgement and damage. The window also includes a dedicated injection port, allowing the local administration of fluids, dyes or drugs with ease (Fig. 1B [3]).

**Figure 1.**
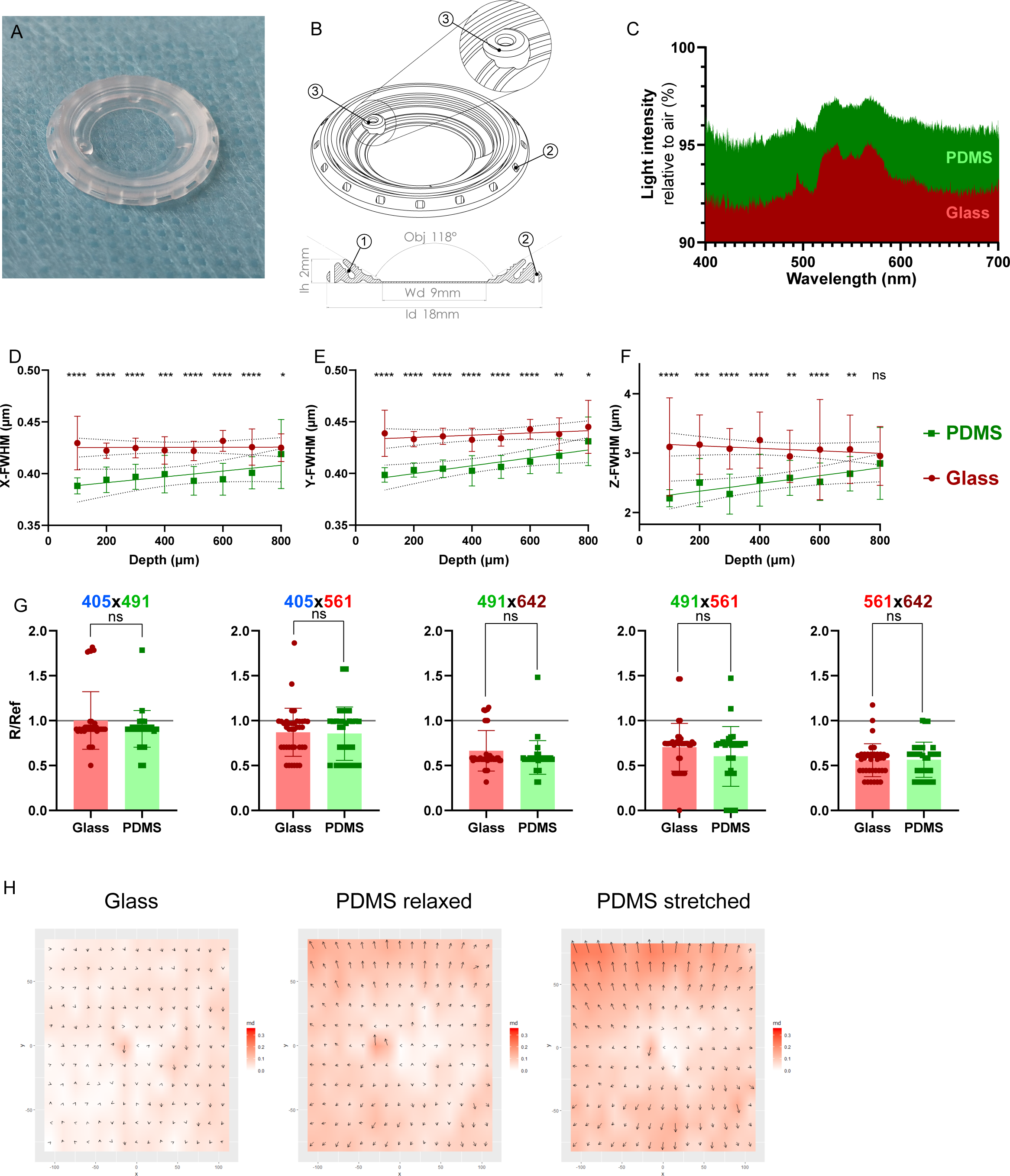
PDMS implantable devices display microscopy-ready optical characteristics. **(A)** Photograph of a PDMS intravital imaging window. (**B)** 3D drawing and cross-sectional view of the window; 18mm in diameter (ld) and 2mm in height (Ih). The device allows imaging with an angled objective (Obj) up to 118° over a working diameter (Wd) of 9mm. At 200mg, the PDMS window is at least 3x lighter than conventional titanium/glass windows. The angle of the skin groove [1] protects the window frame from damage and removal by the mouse. To promote healing and immobilisation of the window, equally spaced holes surround the structure [2]. A port for safe injection [3] is positioned on the side of the window. (**C)** Light intensity measured through glass (red) or PDMS (green) windows relative to air. **(D-F)** Point Spread Function analysis comparing glass (red) and PDMS (green). Graphs display calculated Full Width at Half Maximum (FWHM) values in (**D)** X axis, (**E)** Y axis or (**F)** Z axis over depth. Comparison between glass and PDMS FWHM at each depth was performed using a Mann-Whitney test (n > 20 measurements per section). (**G)** Chromatic aberration analysis of the indicated colour pairs through glass (red) or PDMS (green) windows. Both glass and PDMS in every colour pairs are significantly lower than ratio R/Ref < 1, indicative of co-localisation (p-values < 0.01; One-sample Wilcoxon text). No significant differences were observed between glass and PDMS for all colour pairs (p-values > 0.05, Mann-Whitney test). (**H)** Mapping of the micro-grid deformations arising from glass coverslips, relaxed or stretched PDMS windows compared to the micro-grid alone. Vectors of deformation were calculated on each dot of the grid to generate 2D maps (md = measured deformation norms (μm), vectors represented at 50x). Deformations induced by glass coverslips (0.037±0.019 μm SD), relaxed (0.076±0.031 μm SD) and stretched (0.099±0.043μm SD) PDMS windows were all below the lateral imaging resolution (0.161μm/pixel) and the inherent aberrations of the optical system. P-values: ns = non-significant; * < 0.05; ** < 0.01; *** < 0.001; **** < 0.0001. All source data are provided in **Supplementary Files 2 and 3**.

### Optical properties of PDMS-based intravital imaging windows

The PDMS window is highly transparent, possessing a RI of 1.38±0.05 - in between water (~1.33) and glass (~1.52) - with equivalent light transmittance to glass coverslips of a similar thickness (Fig. 1C, Fig. S1A-B). For decreased light scattering, and thus optimal light penetrance, most IVM studies rely on multiphoton microscopes equipped with pulsed infrared two-photon lasers (two-photon excitation fluorescence microscopy; 2PEF) and water-based objectives for deep-tissue imaging (*3*, *22*, *23*). To compare the resolution of the confocal signal through PDMS with conventional glass coverslips, we performed point spread function (PSF) analyses of 0.2 μm fluorescent beads embedded in agarose (Fig. S1C). The calculated Full Width of Half Maximum (FWHM) of signal intensities (Fig. S1D) obtained via PDMS significantly outperformed corresponding glass measurements in lateral (x, y axis; Fig. 1D-E) and, particularly, axial planes up to 700μm (z axis; Fig. 1F), imaging depths routinely achieved by IVM. Notably, X/Y ratios of the calculated FWHM for PDMS and glass were also comparable, showing similar means and dispersion close to a ratio of 1, indicative of perfect XY squareness (Fig. S1E). This implies that potential subtle deformations of the flexible window do not result in micro-scale geometrical aberrations during imaging. Moreover, using four-color fluorescent beads embedded in agarose (Fig. S1F), we did not observe any differences in the degree of chromatic aberrations induced by glass or PDMS, when both materials displayed co-localization (R/Ref < 1) of specific channel pairs (Fig. 1G). Finally, as macro-scale morphological aberrations arising from the imaging surface are a concern for soft materials, we used a fluorescent stepped micro-grid to assess the deformation generated by PDMS windows in a relaxed or stretched state (Fig. S1G). While morphological aberrations induced by PDMS windows were marginally more pronounced than that induced by glass coverslips, these were negligible compared to the lateral imaging resolution and the inherent anomalies of the optical system (Fig.1H, Fig. S1H). Collectively, these results validate the PDMS membrane as an appropriate optical material for IVM, which is equivalent, and in some cases superior, to glass for water-based immersion imaging.

### Intravital imaging of dynamic, rapidly growing tissues through PDMS imaging windows

Next, we sought to test the long-term maintenance and imaging performance of our PDMS device in biological contexts where rapid tissue growth or anatomical constraints pose significant challenges for IVM using rigid windows: the branching mammary gland during puberty and pregnancy, Patient Derived Xenograft (PDX) tumors and joint muscles. Our optimized surgical protocol (schematically represented in Fig. 2A and described in detail in Methods) facilitates rapid (~5 min) window implantation over superficially located tissues that do not require additional steps for organ exposure. As the device can be folded, the initial incision can be smaller than the circumference of the window for insertion (Fig. S2A-B), and subsequently extended to precisely fit the frame (Fig. 2A). Moreover, pre-positioning the window typically takes less than 3 minutes from the initial incision (Fig. S2A-B), reducing the risk of infection and tissue desiccation prior to the final window-fitting step (Fig. 2A, Fig. S2C). As a result, animals rapidly recover from the short surgical procedure, and comfortably maintain windows for up to 35 days (maximum time tested) (Fig. S2D, Movies S1 and S2). Importantly, the suture-free implantation procedure does not require advanced surgical skills and can be rapidly conducted by non-expert users, a significant improvement over conventional windows. As the design is aimed at limiting constrains on the underlying tissues, the flexible window cannot be fixed for imaging. Thus, to stabilize the device and reduce motion artefacts during IVM, we also developed a custom-made holding system compatible with upright microscope configurations and high numerical aperture ceramic objectives with shallow nose-cones (Fig. S2E-H, Movie S3, Data S1).

**Figure 2.**
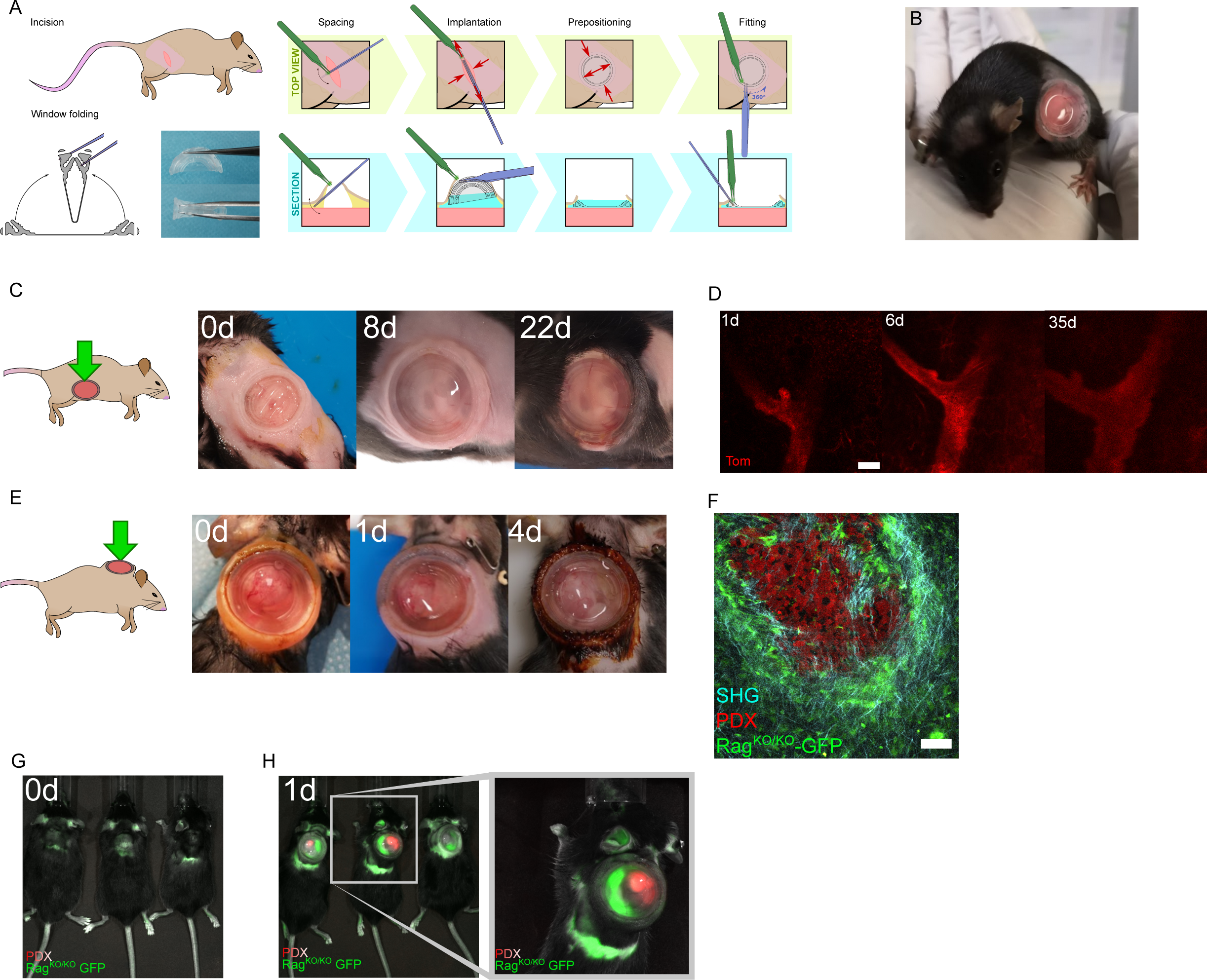
PDMS-based intravital windows allow high-resolution longitudinal imaging *in vivo.* **(A)** Schematic representation of the suture-free window implantation procedure. For details see Methods section. **(B)** Representative photograph of a 6-week-old mouse carrying a newly implanted PDMS window over the abdominal mammary gland. **(C)** Photographs of a PDMS window maintained over the 4^th^mammary gland of a pubertal mouse for the indicated period in days (d). Representative of n > 10 mice implanted at 5-6 weeks of age. **(D)** Longitudinal tissue-scale IVM of a mammary duct expressing tdTomato (Tom) in a *R26^mTmG^* mouse for the indicated period in days (d). Scale bar: 100μm. Maximum tissue depth achieved: 400μm (day 35). **(E)** Photographs of a PDMS intravital imaging window maintained on a growing PDX tumour grafted in the interscapular region of a Rag^KO/KO^-GFP mouse for the indicated period in days (d). Brown skin colouring is due to dermal betadine solution. Representative of n=7 mice. **(F)** IVM of tdTomato-expressing PDX tumour cells (red) engrafted in a Rag^KO/KO^-GFP recipient mouse (host cells in green). Second harmonic generation (SHG) shows fibrillar collagen around the tumour (blue). Scale bar: 100μm**. (G-H),** Near Infrared imaging (NIR-IVIS) of Rag^KO/KO^-GFP mice bearing PDX tumours expressing tdTomato fluorescence before **(G)** or after **(H)** window implantation.

To assess the compatibility of our device with longitudinal 2PEF IVM spanning several weeks, we implanted the window over the 4^th^abdominal mammary gland of pubertal and pregnant R26^mTmG/+^reporter mice (*24*), where all cells are labelled with membrane tdTomato fluorescence (Fig. 2B-D, Fig. S3A-D). Despite substantial increases in body size during puberty, PDMS-windows were well maintained over time, allowing specific tissue regions to be recognized in successive imaging sessions (Fig. 2C-D). Window implantation at late pregnancy also allowed the flexibility of our device to be tested under extreme conditions of skin stretching and rapid body growth (corresponding to a 10% increase over 3 days) (Fig. S3A). In this context, we could visualize lobulo-alveolar development in the pre-lactating mammary gland at high cellular resolution (Fig. S3B-C). Moreover, implanted windows had no detrimental impact on normal parturition nor the lactational competence of the underlying mammary gland (Fig. S3D). Occasionally, IVM post-implantation revealed an accumulation of immune cells underneath the window that obscured underlying epithelial structures (Fig. S3E). This was also reported with conventional glass/titanium imaging windows (*6*, *13*), suggesting that local inflammatory reactions may occur in response to surgery. To mitigate this issue, we performed saline washes prior to imaging via the integrated window injection port (Fig. 1B [3]), which provides a low-risk and non-invasive method for conserving image quality over time (Fig. S3E).

Next, we evaluated the performance of our device over rapidly growing PDXs from Ewing sarcomas engrafted in the interscapular region of immunodeficient NMRI-Nude and Rag^KO/KO-GFP^mice (*25*, *26*). Importantly, PDMS windows implanted over tumors in this location were well-tolerated by both strains (Fig. 2E, Fig. S3F, Movie S4). To enable high-resolution fluorescence IVM of tumors, we generated tdTomato-expressing PDX tumor cells and grafted them in Rag^KO/KO-GFP^host mice. Windows were subsequently implanted over tumors approximately 400 mm^3^in size, which continued to grow rapidly underneath the device at a daily rate of 100-200 mm^3^until reaching their defined endpoint size (Fig. 2E). Despite the anatomical constraints posed by this implantation site, we were able to obtain stable and high-quality IVM images of tumor cells (red), surrounding host stromal cells (green) and the encapsulating tumor collagen network (second harmonic generation, blue) through PDMS windows (Fig. 2F). PDXs are commonly used in pre-clinical studies, which typically rely on Near Infrared (NIR) imaging systems (e.g. IVIS) for monitoring tumor responses over time. However, these systems are poorly compatible with fluorescence imaging and, as expected, standard IVIS imaging failed to detect tdTomato fluorescence in our PDXs models (Fig. 2G). In contrast, a clear and well-defined signal was detected for both green (host cells) and red (tdTomato-expressing tumor cells) fluorescence after PDMS window implantation (Fig 2H). Thus, the application of our PDMS window in this context enables increased reporter sensitivity, in addition to the pre-screening and prioritization of mice for downstream high-resolution confocal microscopy, representing a valuable improvement to canonical NIR studies.

### IVM of muscle regeneration through PDMS imaging windows

Finally, to test other body locations that require high levels of flexibility, we implanted the window over the mouse lower back and thigh muscles (Fig. 3A-B). These anatomical positions are uneven and adjacent to joints, leading to continual deformation of the window and its frame in multiple axes with body movement. While typically problematic with conventional rigid imaging devices - restricting studies to acute or short-term imaging (*27*–*29*) - our implanted PDMS windows were well-maintained at these locations for at least 3 weeks (maximum time examined). To validate the utility of the device for visualizing dynamic biological processes over time, we sought to monitor stem cell activation during injury-induced muscle regeneration in Pax7^CreERT2/+^; R26^mTmG/+^mice (*24*, *30*). Tamoxifen administration in this model induces permanent GFP labelling of Pax7-expressing muscle stem cells and their progeny, allowing their dynamic behaviors in response to muscle injury to be visualized *in situ*. To induce muscle injury in Pax7^CreERT2/+^; R26^mTmG/+^mice, cardiotoxin was administered by intramuscular injection either during the window implantation procedure or after, using the inbuild window injection port to test the feasibility of accurate tissue targeting underneath the device. High-resolution IVM imaging through the PDMS window revealed the movement and division of GFP+ cells at 3 days post-injury (Fig. 3C, Movie S5), a time point at which muscle stem cells are maximally proliferating (*29*, *31*). Importantly, high-resolution images could be acquired through PDMS windows with no observable decline in image quality for at least 18 days after injury (Fig. 3D), allowing the process of muscle regeneration to be visualized in its entirety. Thus, our device is ideally suited to high-resolution longitudinal IVM at the cell and tissue scale of dynamic processes that occur over prolonged timeframes, including in challenging anatomical sites where rigid windows cannot be maintained long-term.

**Figure 3.**
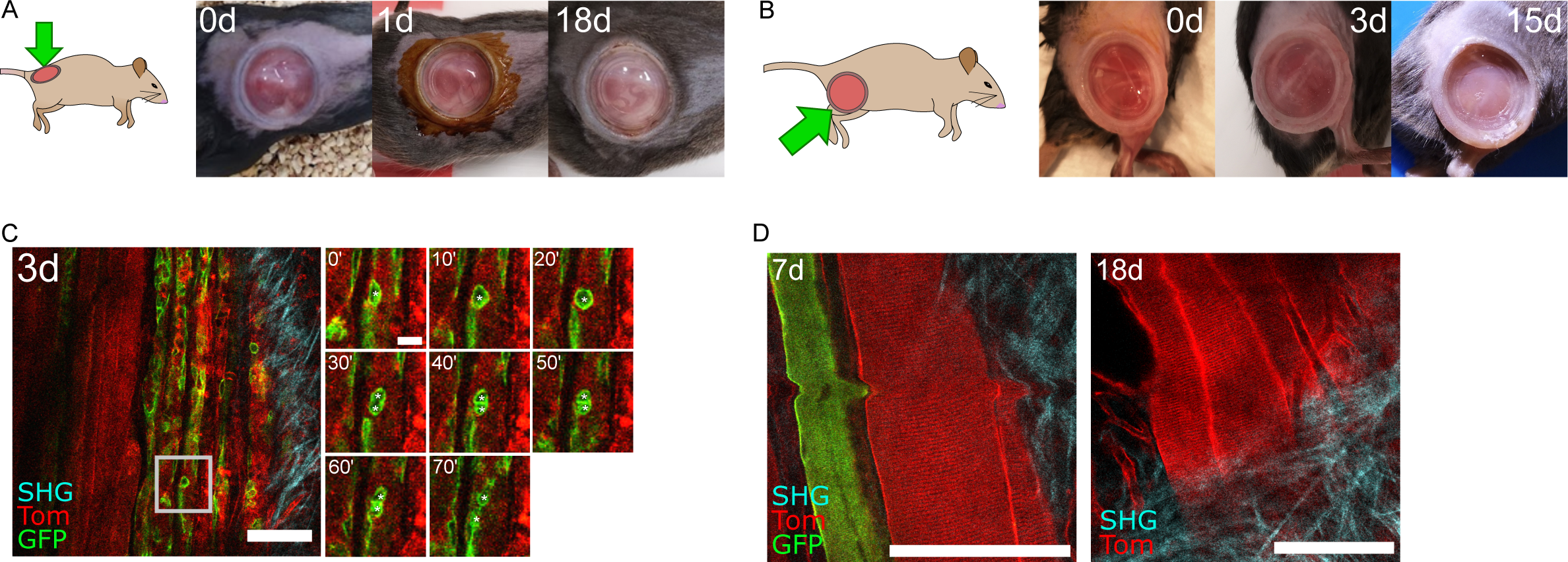
PDMS-based intravital windows allow high-resolution longitudinal imaging of injury induced muscle regeneration *in vivo*. **(A-B)** Representative photographs of a PDMS intravital imaging window maintained on the back **(A)** or thigh **(B)** muscle of a mouse for the indicated period in days (d). Brown skin colouring at day 1 in **(A)** is due to dermal betadine solution. Representative of n = 9 mice implanted on the back or thigh muscles. **(C)** IVM of thigh muscle regeneration through a PDMS imaging window 3 days (3d) after Cardiotoxin-induced muscle injury in *Pax7^CreERT2^;R26^mTmG^* adult mice. All cells were labelled with membrane tdTomato fluorescent protein (Tom, red). Tamoxifen induced Cre-mediated EGFP protein expression (green) in *Pax7*-positive muscle stem cells and their progeny. Second harmonic generation (SHG) imaging shows fibrillar collagen organisation in blue. Time-lapse imaging shows muscle stem cell division and migration (white asterisks) over 70 minutes. Scale bar: 100μm. **(D)** Representative IVM images of muscle fibres 7 and 18 days after implantation in a *Pax7^CreERT2^;R26^mTmG^* mouse, showing resolution at the cellular scale and maintenance of excellent imaging quality over time. Scale bar: 100μm. The small bump in the image at 7 days is due to breathing-induced movement during image acquisition.

## Discussion

Dynamic visualization of cellular processes unfolding inside living animals is essential for an in-depth understanding of normal and pathological organ function. A major challenge thus far has been the ability to study cellular dynamics in vivo, without perturbing tissue physiology. Intravital imaging represents a critical advance in this respect. Nevertheless, the ability to concomitantly expose and preserve tissues for long-term imaging remains a fundamental challenge, one largely inaccessible to most laboratories. The development of a flexible, transparent and seamlessly integrated “second skin” - as afforded by our imaging window - represents a marked advance in methodology. Our suture-less PDMS-based device possesses significant technical, biological and ethical advantages over conventional imaging windows. This includes a portal for *ad hoc* local delivery of substances, in addition to a rapid and straightforward implantation. The lightweight and low-profile design of the window also ensures minimal impact on normal animal behaviors, in addition to facilitating group housing and the safe inclusion of environmental enrichment, an important improvement on current welfare protocols. Moreover, implantation of PDMS imaging windows lead to substantial improvements in the performance of NIR-imaging systems for *in vivo* fluorescence detection. Importantly, this window is also compatible with other imaging technologies, including MRI and echography. Thus, our device is ideally suited for developmental, pathological or pharmacokinetic studies across multiple imaging scales.

While particularly suited to superficial tissues in dynamic and flexible locations, the versatility of PDMS makes our model readily adaptable for internal organ imaging, by incorporating additional surgical steps for organ exposure or silicone-based glues. The suppleness of the device, however, renders it less suitable for implantation over rigid tissue sites, such as for brain or thoracic organ imaging that demand more specialist solutions (*10*, *18*, *19*). Nonetheless, silicone mass or surface modifications, in addition to drug embedding, can be used to alter the window’s mechanical, adhesive or bio-activity properties for enhanced healing, tissue anchoring, permeability and resistance to biological/chemical agents. As substrate rigidity plays important roles in tissue development and tumorigenesis (*32*), the more physiological stiffness of PDMS - alongside the ability to tune its mechanical properties for specific research needs - is also an important advantage over conventional glass coverslip models. Moreover, in light of exponential progress in microfluidic devices using the same production methods, integrating micro-patterns, electrodes or surface modifications into the window design for live sensing or interaction with the underlying tissue represents a realistic future advance. Finally, in contrast to currently available solutions, plastic engineered devices such as ours are scalable, allowing single-use and widespread distribution at low cost for basic and preclinical research.

## Materials and Methods

### Animals

All studies and procedures involving animals were in strict accordance with the recommendations of the European Community Directive (2010/63/UE) for the protection of vertebrate animals used for experimental and other scientific purposes. The project was specifically approved by the ethics committees of Institut Curie CEEA-IC #118 and Institut Pasteur (reference #2015-0008), and approved by the French Ministry of Research (authorization references #0424003, APAFIS #13310-2018020112578233-v1, APAFIS#6354-20160809l2028839v4, APAFIS#11206-2017090816044613-v2). We comply with internationally established principles of replacement, reduction and refinement in accordance with the Guide for the Care and Use of Laboratory Animals (NRC 2011). Husbandry, supply of animals, as well as mouse maintenance and care in the Animal Facility of Institut Curie (facility license #C75–05–18) before and during experiments fully satisfied the animals’ needs and welfare. Suffering of the animals was kept to a minimum.

The following mouse lines were used in this work: *R26^mTmG^* mice(*24*); NMRI-Foxn1 nu/nu (purchased from Janvier Labs*); Pax7^CreERT2^* mice(*30*), Rag^KO/KO^-GFP mice (B6(Cg)-*Rag2^tm1.1Cgn^*/J;Tg(UBC-GFP)(*25*, *26*)). All experiments were performed in 4-8-week-old female mice.

### Prototyping

3D conception of the window, mold and holder was performed using Solidworks 2018 (Dassault Systems). 3D printing was performed using a home-made system (FDM) and MultiJetFusion (HP) using various materials, predominantly generic Acrylonitrile Butadiene Styrene (FDM), Polyamide and Polypropylene (PP) (fusion). We recommend PP-fusion for higher printing resolution, temperature and hygroscopic stability.

### Window production

Biocompatible transparent silicone (Elkem) components (A and B) were mixed at a 1:1 ratio (A:B) for 5 min per 50g of mix. The mix was then degassed for 5 min and 10 min (<10mbar) with a vacuum release in between to maximize air removal. Air removal was completed by centrifuging the mix at 1000g for 5 min. The degassed silicone mix was kept for a maximum of 24h at 4°C to prevent spontaneous polymerization. Windows were produced using a custom 3-stage single footprint compression mold maintained at 100°C throughout molding. Approximately 1ml of silicone was placed on the footprint prior to closing the mold and applying compression at 5 tons for 10 min. Polymerized windows were then post-cured for at least 24h at 60°C, and subsequently washed in 100° ethanol before autoclaving.

### Optical characterization of PDMS windows

Optical, chromatic and morphological aberrations depend on the optical path, which includes the objective and microscope core. Therefore, all assays were performed using a single microscope setup, and water as the imaging medium. All measurements were acquired using a 40x/1NA Water DIC PL APO VIS-IR objective on an upright spinning disk microscope (CSU-X1 scan-head from Yokogaw; Carl Zeiss, Roper Scientific France), equipped with a CoolSnap HQ2 CCD camera (Photometrics) and Metamorph software.

#### Optical aberrations (PSF)

To measure the Point Spread Function (PSF) representing optical aberrations arising from refractive index changes, we prepared 2% agarose gels containing 1:200 fluorescein and 1:400 diluted FluoSpheres™ carboxylate-modified microspheres (0.2μm diameter). For both glass and PDMS windows, we acquired 8 stacks of 100 μm with a step size of 0.2μm at increasing depths from the window/gel interface (Z0, defined using the background signal in the fluorescein channel). PSF was calculated using the MetroloJ plugin(*33*) in FIJI (ImageJ v1.53) on at least 20 beads per section.

#### Chromatic aberrations

To measure the chromatic aberrations representing optical aberrations induced by wavelength specific refractive index changes, we prepared 2% agarose gels containing TetraSpeck™ Microspheres (4μm diameter) diluted at 1:400. Data was acquired over 200μm from the first bead detected in Z for both glass and PDMS windows. Laser intensities and exposures were set to obtain similar intensity range (12-bits). R and Rref were calculated using the MetroloJ plugin(*33*) in FIJI (ImageJ v1.53) on at least 30 beads per condition.

#### Morphological aberrations

To measure morphological aberrations that may arise from an uneven window surface, we imaged a microgrid (Argolight SLG-075, Pattern B) using a laser excitation wavelength of 405nm through either water only (uncovered), glass, relaxed PDMS or stretched PDMS (wavy) window in duplicates. Dot grids were segmented using FIJI v1.53 (Threshold>Analyse Particles) and dot coordinates extracted. Using a central motif (cross 0;0) and the top left dot (−105; −75), we corrected the rotation of the grid under the different materials and compared the corrected coordinates of each dot to either “medium only” (measured) or the theoretical (15×15μm) Pattern B matrix. Graphical deformation maps were generated using ggplot2 of R(*34*).

### Window implantation

Before surgery, all instruments were sterilized by autoclaving or heat sterilization. PDMS windows were sterilized by autoclaving to avoid darkening. To ensure sterility throughout the procedure, mouse preparation and handling, adjustment of anesthesia flow rates and similar tasks were performed by an assistant, limiting the surgeon to only contacting sterile surgical tools and the surgical field. The physiological body temperature of the anaesthetized mouse was maintained throughout the procedure using a heated induction chamber and heated surgical station.

#### Preparation (performed by the surgical assistant)

### Pre-operative care

prior to anesthesia, mice were administered 125mg/kg of NOROCLAV® (140mg/ml amoxicillin; 35mg/ml clavulanic acid) and an analgesic cocktail (0.1 mg/kg Buprenorphine (BUPRECARE®) and 5 mg/kg of Carprofen (RIMADYL®)) by sub-cutaneous (SC) injection. Analgesics were administered into the neck skin fold for all experiments apart from implantation in PDX models, where they were administered in the rear leg skin fold.

### Surgical field preparation

Mice were anaesthetized in an induction chamber using 4% isoflurane (ISOFLURIN®). Once anaesthetized, mice were transferred onto a heat-pad, and anesthesia maintained using an isoflurane concentration of 1.5-2%. Ophthalmic ointment was applied to eyes to prevent corneal drying. At the planned implantation site, mouse hair was trimmed using an electronic pet clipper (Aesculap® Exacta). Residual hairs were removed using depilatory cream for a maximum of 2 minutes to minimize the risk of skin damage and drying. Stray hairs and cream were removed using BETADINE Scrub 4%, prior to a soap rinse with sterile water. The shaved skin and surrounding areas were subsequently disinfected using dermal BETADINE 10%. Finally, the prepared surgical area was covered by pre-cut sterile film (Tegardem™).

#### Window implantation (performed by the surgeon, procedure illustrated in Fig. 2A)

We recommend the following surgical tools to limit skin and tissue damage: Graefe extra fine straight 1×2 teeth forceps (for skin and membranes - FST #11153-10), flat blunt forceps (for the window and tissue), serrated blunt forceps (for the window), fine, sharp straight scissors (for skin and membranes - FST #14568-09) and Strabismus blunt straight scissors (for blunt dissecting the subcutaneous pocket FST #14574-09).

To implant the PDSM-window (Ø18mm) in the prepared surgical area, a 10mm incision was made in the skin overlying the tissue of interest (corresponding to approximately 15mm when stretched). Next, using blunt scissors, the skin was carefully detached from underlying tissues approximately 5-10mm around the incision to generate sufficient space for the implanted window. Throughout the procedure, pre-warmed saline solution (sterile 0.9% NaCl) was used to prevent tissue dehydration. Prior to positioning the window, the incision site was filled with pre-warmed saline solution to minimize the risk of introducing air underneath the window during implantation. Using thin forceps with blunt and flat edges, the PDMS-window was folded in half and placed into the incision pocket where it was unfolded under the skin. Typically, the procedure from the first incision to pre-positioning of the window under the skin was performed in under 3 min (**Fig. S2a-b**). To fit the window in place, the skin edges were positioned into the window groove using two forceps that alternate between holding the upper frame and sliding the skin around the window (similar to the technique for adjusting a tire on a rim). During this step, the incision size can be adjusted, if required, to tightly seal the skin around the window. Experienced users are capable of performing the entire surgical procedure in ~ 5 min.

### Post-operative care

After surgery, mice were placed in a pre-warmed cage (on a heating pad) to recover from the anesthesia. Mice were closely monitored for signs of pain, discomfort and infection after window implantation. The second and last NOROCLAV antibiotic dose was provided by SC injection the day after surgery. The skin surrounding the window was cleaned using Dermic BETADINE 1% every 2-4 days in order to prevent infection. Ibuprofen was administered in drinking water (0.4 mg/ml) for 7 days post-surgery to minimize inflammation and pain. If signs of pain are observed, post-operative analgesics (0.1 mg/kg Buprenorphine (BUPRECARE®) and 5 mg/kg of Carprofen (RIMADYL®)) can be administered by SC injection the day after surgery.

Mice bearing windows were group-housed where possible after surgery, and provided with cage enrichment suitable for post-surgery recovery, including nesting material, housing and chew sticks. Recovery food (DietGel® *Recovery*, ClearH20) was provided in the cage to limit mechanical stress from reaching the feeder. Singly-housed mice without sufficient environmental enrichment were more likely to damage the device during the first 1-2 days after surgery. Overall, younger animals were more prone to skin damage arising from the procedure, which may be due to the window/animal size ratio and/or increased activity at earlier ages. Cage cleanliness did not appear to affect the risk of implant infection. Typically, mice exhibited minor weight loss the day after surgery (−4.20%±4.74%, within ethical limits), which were restored by day 2 (+2.93%±3.18%).

### Handling window-bearing mice

When the implantation is performed correctly with stringent asepsis, no mouse should spontaneously lose their implanted windows. Windows are at a risk of displacement, however, in the first few days after surgery when handling mice using conventional restraining techniques. Thus, we recommend adapting handling techniques when assessing mice in the days immediately after window implantation, and consider the use of light gas anesthesia where possible e.g. when performing procedures such as IP injection.

### Troubleshooting guide for the implantation procedure

#### [A] Air is trapped under the window during implantation

Prepare a 1ml syringe with sterile, pre-warmed 0.9% NaCl and a 30G needle. Position the mouse so that the injection port is facing upward and disinfect it with 70% ethanol. Using fine forceps, hold the injection site and inject 200-500μl of saline solution underneath the window: the air bubble(s) will move close to the injection site. Slowly aspirate using the syringe to remove air and as much of the saline solution as possible. Dry the injection port. Wait for the saline solution to be absorbed prior to imaging.

#### [B] Air appears under the window after implantation

During surgery, take care to make a straight incision. Irregular and ragged incisions increase the risk of air leakage. Check the window integrity: if the window is perforated, it must be discarded. Check carefully the skin in the groove for breakages as small cuts can cause air leakiness. If a small break or nick (1-2mm max) is observed, place a small amount of glue on the groove to seal the leakage and proceed to [A]. The next day, check carefully for the absence of air bubbles.

#### [C] Presence of small amount of fluid (exudate) and immune cells between the tissue and the window

This is common for a few days post-surgery and should disappear after 2-3 days. To minimize this risk, review the surgery procedure to ensure stringent asepsis, check/change the ibuprofen in the drinking water and ensure that food is accessible to limit mechanical stress. If the exudate precludes imaging, administer pre-warmed saline through the injection port to flush underneath the window. (see [A], **Fig. S3E**).

#### [D] Hair underneath the window

Review the preparation step. Trim a larger area if necessary. A spray plaster can be used prior to incision to trap residual stray hairs.

#### [E] Skin becomes thinner or dry around the window

If this issue occurs within the first 7 days from implantation: reduce depilatory cream incubation and/or change the brand. Use 1×2 teeth forceps to reduce skin damage. Limit damage to vasculature when performing blunt dissection to detach the skin. This issue may appear after long-term maintenance (> 2 weeks), but in our experience it did not affect imaging (no infection, no air bubbles) when post-surgery care was properly performed.

#### [F] Skin or tissue become necrotic

Immediately terminate the experiment. This should never happen; review the surgery procedure (including asepsis, tissue drying and damage to vasculature).

### Intravital imaging through PDMS-windows

Imaging on the day of implantation is not recommended to avoid additional stress to the surgery area and over-constraining the window. Residual physiological saline under the window immediately after implantation can also affect imaging depth. Instead, perform imaging the next day (day 1) at the earliest. While the mouse is anaesthetized, administer post-operative antibiotics by SC injection. Carefully inspect the device and clean the surgical area as described above. Mice are gas-anaesthetized using 4% isoflurane (ISOFLURIN®) initially and maintained using 1.5-2.5% isoflurane for short term imaging. When performing time-lapse imaging spanning several hours, reduce the isoflurane concentration to between 0.8%-1.2%. These anesthesia levels are optimal for long-term maintenance of mice in a nonresponsive state with a slow, constant and non-forced breathing pattern. Irregular and abnormal breathing patterns are associated with persistent anesthesia greater than 1.5%, which perturbs imaging (increases motion-induced image deformations) and can decrease survival times (*35*). Fast breathing and reflex responses indicate insufficient anesthesia, which also disturbs imaging and increases the risk of the mouse awakening during imaging.

### Before imaging

To maintain animal hydration for short-term imaging (< 2-3 hours), administer 250μl/10g of saline solution by sub-cutaneous injection after anesthesia induction. In experiments exceeding 3 h, mouse hydration can be maintained during imaging using a subcutaneous or intraperitoneal infusion of glucose and electrolytes (~ 50–100 μl/hour) via an indwelling line. Our custom-made holder, adapted for upright microscopes (ready to print STL files provided in **Data S1**), permits regional body immobilization through compression. A rigid conical structure (containing immersion medium, usually water) presses on the edge of the window, allowing the tissue and window to be exposed and stretched beneath the objective as the mouse rests on a foam bed, which preserves breathing while cushioning movements. The mouse is positioned on the imaging holder and gently pushed on the static stage, centering the window in the conic aperture. The mouse should be tightly constrained; the foam absorbs excessive compression. The mouse is then observed for 5 min to check breathing and the stability of the anesthesia. Prior to placing the holder onto the microscope stage, the conic aperture is filled with 2-3ml of pure water. The holder may be adapted for inverted microscope configurations, using gel-based immersion medium instead of water for imaging. **Critical: do not use oil-based imaging medium with PDMS windows**.

### During imaging

Find structures of interest using brightfield or fluorescence with the oculars. During scanning, optimize laser power and exposure times to minimize photo-toxicity and pixel saturation.

### After imaging

The imaging medium must be removed from the conic aperture using a paper towel. The mouse is released from the holder and placed on a heated pad (set at 38°C) for recovery. During this time, the window is cleaned using a paper towel and 70% ethanol and the surrounding skin disinfected with Dermic Betadine 1%.

### Troubleshooting guide for imaging procedures

#### [G] The window cannot be correctly positioned on the conic aperture

Use softer foam in the holder that allows fitting to any position.

#### [H] The imaging medium (water) leaks

The conic aperture is not centered on the window. Reposition it or use a gel-based imaging medium.

#### [I] The holder disturbs breathing

Reduce the isoflurane dose. Reduce the compression on the animal. Sculpt the foam, ensuring that the mouse’s airway is not restricted. Chisel the foam to modify the mouse position (a horizontal plane may not be optimal for the tissue of interest).

#### [J] Poor visibility during imaging

Exudate accumulation and/or epithelioid “membrane” formation beneath the window (which can develop with long-term maintenance) can interfere with imaging. This has also been observed with glass/titanium windows (*6*, *13*), and is likely due to inflammatory reactions in response to surgery. To minimize this risk, review the surgical procedure, taking care to limit tissue damage and maintain stringent asepsis throughout. While the epithelioid membrane-like structure affects tissue imaging depths, overall, it has no detrimental impact on image quality. However, when coupled with immune cells and/or fibrosis, it can preclude imaging. To alleviate this issue, pre-warmed saline can be administered via the injection port to flush underneath the window (following the protocol in [A], **Fig. S3E**). Alternatively, reposition the window on a different area of the tissue of interest, or terminate the experiment.

### Intravital microscopy

Animals were anaesthetized and positioned for imaging using the custom-made holder as described above. Intravital imaging was performed on an upright Nikon A1R MP multiphoton confocal microscope equipped with a pulsed Spectra-Physics Insight DeepSee laser (680-1300 nm, 120 fs pulses with auto-alignment), Luigs&Neumann XY motorised stage and four GaAsP non-descanned detectors with SP492, BP 525/50, BP 575/50 and BP 629/56 filter sets. The microscope was surrounded by a heated dark box maintained at 38°C. All images were acquired using 16x NA 0.8 or 25x NA 1.1 PlanApo LambdaS water objectives. An excitation wavelength of 960 nm was used for GFP and TdTomato, in addition to second harmonic generation (SHG) imaging of collagen.

### Image processing and visualization

Time-lapse acquisitions were corrected for movements and drifts using the “Correct 3D drift” plugin(*36*) in FIJI (ImageJ v1.53) using the SHG channel as a reference.

### Generation of tdTomato-expressing PDX tumor cells

Ewing Sarcoma PDX tumors (IC-pPDX-87) were surgically removed and enzymatically dissociated to single cell level using the protocol described in Stewart *et al* (*37*). Cells were resuspended (2.25 x 10^7^cells/1.5mL) in DMEM/F12 supplemented with 1% penicillin-streptomycin and 2% B27 (all from ThermoFisher) and transduced overnight with a TdTomato-lentivirus in a 2ml tube using an Intelli-Mixer RM-2L (Dutcher), program F1, 5rpm, in a 37°C incubator with 5% CO_2_. TdTomato-lentivirus was generated by replacing the GFP sequence with tandem Tomato in the Lenti-sgRNA-GFP (LRG) plasmid (Addgene plasmid # 65656; http://n2t.net/addgene:65656, a kind gift from Christopher Vakoc). The next day, cells were spun at 500g for 10 min. For one injection, 1 x 10^7^cells were resuspended in 50μL of media and kept on ice. 50μL of Matrigel (Corning, Ref 354234) was added to the cell suspension prior to injection into the interscapular fat tissue using an insulin syringe (BD ref 324891).

### Muscle regeneration studies

Muscle injury was performed in *Pax7^CreERT2^;R26^mTmG^* adult mice at the time of PDMS-imaging window implantation by intramuscular injection of 50 μl of Cardiotoxin (10 μM, Latoxan, L8102) diluted in 0.9% NaCl. Cardiotoxin was administered either before window positioning, or after through the injection port. Mice received 3 mg of tamoxifen free base (Euromedex) by intraperitoneal injection two days prior to intravital imaging.

### IVIS imaging

NIR fluorescence imaging was performed with a highly sensitive, CCD camera mounted in a light-tight specimen box (In Vivo Imaging System – IVIS 50; Living Image Version: 4.3.1.0.15880). Animals were anaesthetized in a warm induction chamber using 4% isoflurane, and subsequently placed onto a warmed stage inside the camera box and supplied with 1.5-2% isoflurane to maintain anesthesia during imaging. The emitted fluorescence signal was detected by the IVIS camera system, integrated, digitized, and displayed. Excitation filters for Tomato and GFP fluorescence were 535 (DsRed filter, Position 3) and 465 (GFP filter, Position 2) respectively. Exposure times and pixel binning were auto-optimized for each fluorescence channel to minimize over-exposure and normalized for comparison using the formula *Normalised_Intensity = (Measured_Intensity / Exposure_Time) / Binning_Factor².* Raw parameters are provided in **Data S2**.

## Acknowledgments

We particularly acknowledge Philippe Jacquemin (Lycée Marcelin Berthelot, Questembert – plastic engineering section) for his help on window and mould design, initial prototyping and for teaching us the basics in polymer properties and processing. We especially thank Lucie Sengmanivong, Marie Irondelle and Olivier Leroy from the Cell and Tissue Imaging (PICT-IBiSA) facility and the Nikon Imaging Centre, Institut Curie, member of the French National Research Infrastructure France-BioImaging (ANR10-INBS-04). We are also grateful to Raphael Margueron (Institut Curie) for advice and constructive discussions. We wish to acknowledge Olivier Delattre and Sakina Zaidi (Institut Curie) for providing the PDX tumours, in addition to the Institut Curie In Vivo Experimental Facility for help in the maintenance and care of the mouse colony.

## Funding

This work was supported by grants to S.F., B.L-L and G.J. from Paris Sciences et Lettres (PSL* Research University), Institut Carnot, the French National Research Agency (ANR - grant # ANR-15-CE13-0013-01), the Canceropole Ile-de-France (grant # 2015-2-APD-01-ICR-1), the Ligue contre la cancer (grant #RS19/75-101) and by the Labex DEEP ANR-Number 11-LBX-0044, as well as the "FRM Equipes" EQU201903007821 and the FSER (Fondation Schlumberger pour l’éducation et la recherche) FSER20200211117. The PICT-IBiSA facility is supported by the Fondation pour la Recherche Médicale (FRM # DGE20111123020), the Canceropole Ile-de-France (# 2012-2-EML-04-IC-1), InCA (Cancer National Institute, # 2011-1-LABEL-IC-4). D.S and A.B. are supported by the Institut Curie-SIRIC (Site de Recherche Intégrée en Cancérologie, INCa-DGOS-Inserm_12554; ITMO Cancer AVIESAN). B.L-L acknowledges support from the Academy of Medical Sciences, Wellcome Trust and the University of Bristol. S.T acknowledges funding support from the Institut Pasteur, Agence Nationale de la Recherche (Laboratoire d’Excellence Revive, Investissement d’Avenir; ANR-10-LABX-73), M.B.D. was supported by the Laboratoire d’Excellence Revive and La Ligue Contre le Cancer.

## Author contributions

G.J., S.F., and B.L-L., conceived and administered the study. G.J., M.B.D., V.D-M., D.S., S.T., S.F. and B.L-L., conceived and designed experiments. G.J., M.B.D., S.D., A.B. and B.L-L. performed experiments. G.J., S.F. and B.L-L., wrote the manuscript. All authors reviewed and approved the manuscript.

## Competing interests

The PDMS imaging window described in this publication is patented under the number #EP3656349A1 by Institut Curie, where G.J., B.L-L. and S.F. are named inventors.

## Data and materials availability

Data supporting the findings of this study are available within the article. Source data are available in the manuscript files. Requests for further information should be directed to corresponding authors.

## Notes

### Competing Interest Statement

The PDMS imaging window described in this manuscript is patented under the number #EP3656349A1 by Institut Curie, where G.J., B.L-L. and S.F. are named inventors.

