## Supplementary Materials for "Longitudinal high-resolution imaging through a flexible intravital imaging window"

### EXTENDED FIGURE 1

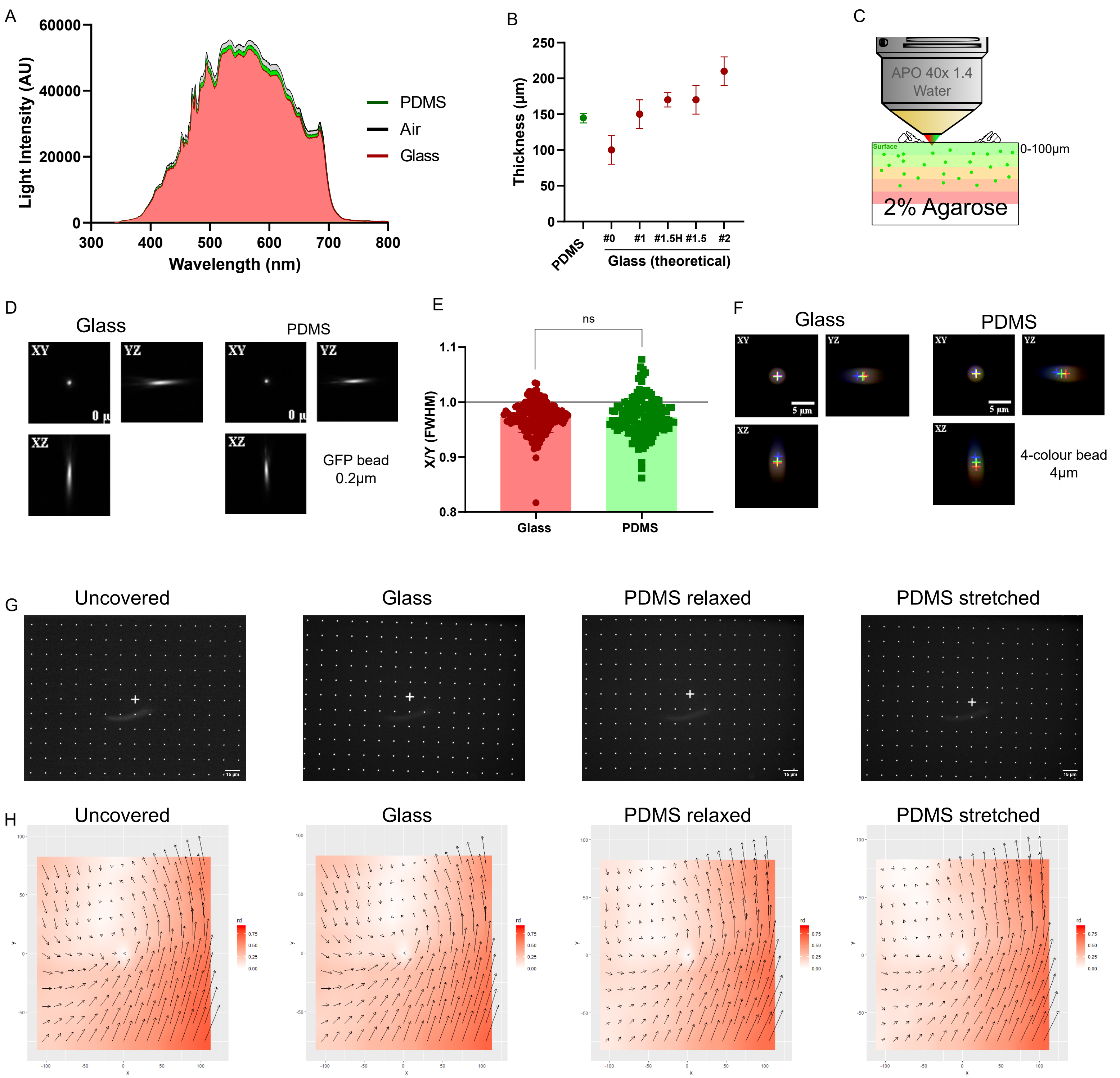

**Fig. S1. Related to Figure 1.**

**(A)** Profile of absolute light intensity measured in air (grey), glass (red) and PDMS (green) for wavelengths spanning 300nm to 800nm. **(B)** Experimental measures of the PDMS window thickness (n=10, in green) and theoretical thickness of glass coverslips of different tolerances (ISO8255, in red). **(C)** Schematic of the experimental setup for Point Spread Function (PSF) measurements. **(D)** Example PSF profiles for a 0.2µm diameter GFP bead through glass or PDMS windows. **(E)** XY squareness (X/Y ratio) of the FWHM of 0.2µm diameter GFP beads from 0 to 800µm of depth through glass (red) or PDMS (green), reflecting micro-geometrical deformations. **(F)** Examples of co-localization profiles for a 4µm diameter four-colored bead through glass or PDMS windows. **(G)** Representative pictures of micro-grids acquired by imaging: medium only (uncovered), glass and relaxed or stretched PDMS windows. **(H)** Mapping of micro-grid deformations acquired by imaging: medium only (uncovered), glass, relaxed or stretched PDMS windows compared to a theoretical 15µm step micro-grid showing the geometrical/spherical aberrations associated with the optical system (rd = real deformation norms, unit in µm, vectors represented at 50x). All source data are provided in **Supplementary Files 2 and 3.**

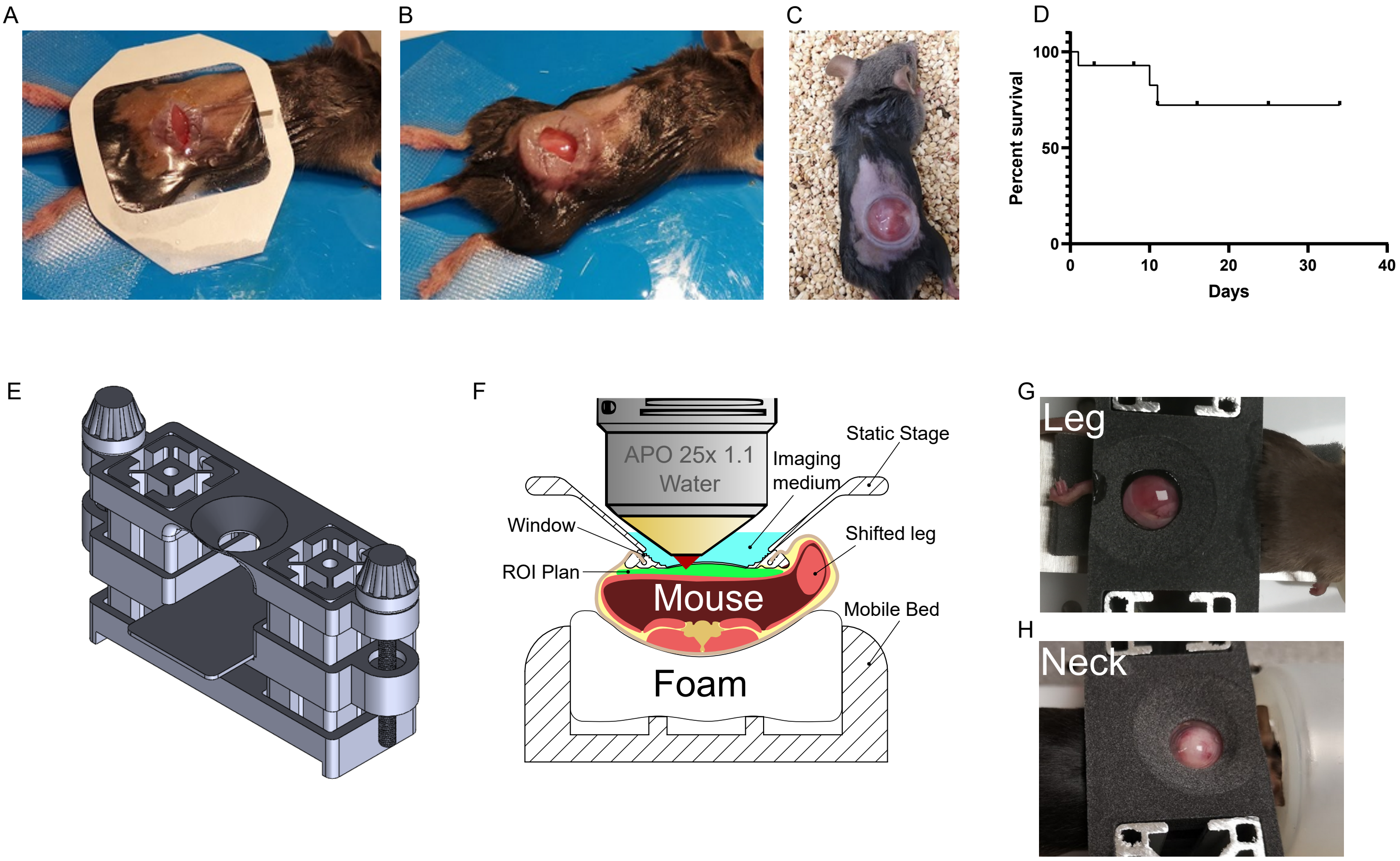
A
B
C
D
Percent survival
100
50
0
0
10
20
30
40
Days
E
F
APO 25x 1.1
Water
Static Stage
Imaging
medium
Window
Shifted leg
ROI Plan
Mouse
Mobile Bed
Foam
G
Leg
H
Neck

**Fig. S2. Related to Figure 2.**

**(A)** Photograph showing the skin incision prior to window implantation on the back muscle. The operated area is protected by a sterile adhesive plaster (Tegaderm$^{TM}$) ensuring aseptic conditions. **(B)** Photograph of the pre-positioned window in the skin pocket. The window is entirely under the skin, protecting the underlying tissues from contamination. **(C)** Mouse recovering from the anesthesia immediately after window fitting. **(D)** Survival curve of mice carrying PDMS windows (n=15). Windows were displaced in three mice, two due to handling, and one due to an infection arising from poor aseptic technique during surgery. We recommend that users minimize restraining mice using conventional handling techniques, particularly in the days following implantation. No spontaneous window loss or clinical signs of distress were observed in > 26 mice. **(E-H),** Three-dimensional rendering of the assembled, custom-made holder **(E)** and schematic diagram of the system during imaging, highlighting its loco-regional compression function **(F).** The holder is adapted for upright microscope configurations and is fully compatible for imaging body sites from the rear leg **(G)** to the neck **(H).** The design can also be adapted for inverted microscope configurations. This holder substantially reduced motion artifacts during two-photon *in vivo* imaging, eliminating the need for gated acquisitions and advanced post-imaging processing methods. The holder also ensured reproducible window positioning in successive imaging sessions. Holder design is included in **Supplementary File 1**.

EXTENDED FIGURE 3

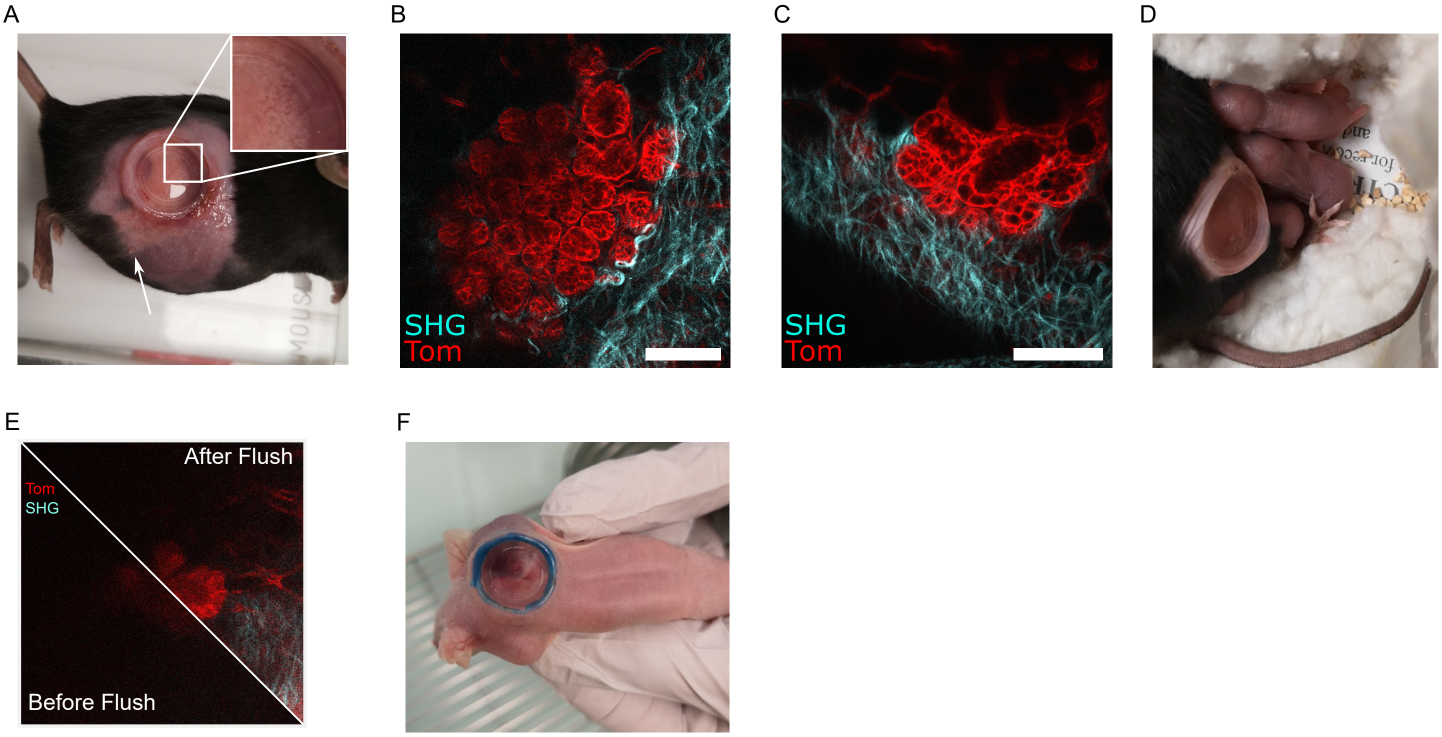

**Fig. S3. Related to Figure 2.**

**(A)** Photograph of a PDMS window placed onto the 4th mammary gland of a $R26^{mTmG}$ mouse at late pregnancy (P16.5) and maintained until parturition (3 days later), despite a 10% increase in mouse body weight during this time, and 25% loss after delivery (n=4). Pregnancy-induced changes to mammary gland architecture were noticeable by eye (inset in a). The white arrow points to the 4th mammary gland nipple, which remained intact and accessible after window implantation. (**B-C)** High-resolution IVM through the PDMS window of developing lobulo-alveolar structures in the mammary gland of a pregnant $R26^{mTmG}$ mouse, labelled with tdTomato red fluorescence (Tom). SHG shows the organization of fibrillar collagen in blue. Scale bar: 100µm. (**D)** Window implantation during pregnancy had no detrimental impact on parturition (11 pups were born) nor on the lactational competence of the underlying mammary gland. **(E)** Representative IVM image of the partial recovery of image quality after flushing immune cells using saline solution administered through the window injection port. Example shows the same mammary epithelial structure labelled with tdTomato fluorescence (red, Tom), before and after the procedure. SHG shows the organization of fibrillar collagen (blue). **(F)** Representative photograph of a NMRI-Nude mouse carrying a PDMS window on a neck-implanted PDX tumor ~3 weeks after window implantation. Blue coloring of the outer frame of the window was used to aid its positioning during implantation.

**Movie file legends:**

**Movie S1**: video showing mice recovering immediately after muscle injury and window implantation. One mouse has completely recovered from the surgery, while the newly implanted mouse (with the window on the back) is still recovering from the anesthesia. Animal movements are slightly impaired due to the applied muscle injury.

**Movie S2**: video showing mice 3 days after muscle injury and window implantation. Window-bearing mice behave normally with no detectable impairment in mobility, or any clinical signs of distress arising from the procedure.

**Movie S3:** video showing the stability of the window using the custom-made holding system. The isoflurane dose is adjusted to obtain stable and normal breathing for 5 min before imaging.

**Movie S4**: video showing normal behavior of NMRI-Nude PDX mice 7 days after window implantation. The elasticity of the hairless Nude mouse skin does not affect the maintenance of the flexible PDMS window.

**Movie S5**: video showing thigh muscle regeneration in $Pax7^{CreERT2};R26^{mTmG}$ mice (selected still images are presented in **Fig. 3C**).

**Data file legends:**

**Data S1:** imaging holder design. 7zip archive containing the 4 parts (STL format) of the holder and one example of inox bed shape (DXF format). "StaticStage^Stage4" part was 3D printed in water-resistant and rigid material (MultiJet-Fusion PP) for better stabilization. Parts are designed for assembly on Rexroth-like 30 aluminum profiles using appropriate nuts. Assembly screws are DIN 912 M5x16. Motion screws capped by "Button^Stage4" parts are DIN 931 M5x60.

**Data S2:** source data file for Fig.1 and Extended Fig.1.

**Data S3:** source data file for Fig.1h and Extended Fig.1h.
